# Gene supplementation in Vanishing White Matter mice ameliorates the disease

**DOI:** 10.64898/2025.12.05.692143

**Authors:** Anne E.J. Hillen, Robbert Zalm, Marjo S. van der Knaap, Vivi M. Heine

## Abstract

Vanishing White Matter is a leukodystrophy associated with neurological decline, including motor deficits, and premature death. It is caused by mutations in the genes encoding the subunits of eukaryotic Translation Initiation Factor 2B (eIF2B), leading to constitutive activation of the Integrated Stress Response, especially in astrocytes. Brain pathology shows immature and dysfunctional glia in the central nervous system. No curative treatment options are currently available for Vanishing White Matter patients. To investigate a gene therapy approach, we supplemented *Eif2b5*-mutant Vanishing White Matter mice with the wild-type *Eif2b5* gene sequence. We tested lentiviral vectors with different promoters to target astrocytes or cells with heightened Integrated Stress Response activity. The regenerative effects of intracerebroventricular injection in neonates were tested in adulthood. Motor skills and pathology in the central nervous system were improved when wild-type *Eif2b5* delivery was targeted towards astrocytes. No negative side-effects of overexpression were observed in any of the groups. In conclusion, gene supplementation in astrocytes is effective in alleviating disease severity, showing promise for gene therapy for Vanishing White Matter patients.

## Introduction

Vanishing White Matter (VWM) disease is a leukodystrophy that most often has its onset in early childhood. The disease is characterized by neurological deterioration, with progressive ataxia, spasticity, epilepsy, and cognitive decline.^1^ The age of onset of VWM is correlated to disease severity, with 60% of cases having a disease onset before 4 years of age.^2^ Stressors, such as minor head trauma and febrile infection, can accelerate deterioration.^3,4^ At present no treatment options are available, and VWM patients often suffer an early death.^2^

Several other leukodystrophies, caused by recessive pathogenic variants in various genes, have shown successful alleviation of disease symptoms following supplementation with the wild-type (wt) gene.^5–8^ Many gene therapies using this approach target a broad range of cells by ubiquitous promoters such as cytomegalovirus (CMV) or phosphoglycerate kinase 1 (PGK).^5,8–10^ Considering the monogenic nature of VWM, gene therapy may be a curative treatment option: VWM is caused by autosomal recessive mutations in genes (*EIF2B1* to *EIF2B5*) encoding the subunits α to ε of eukaryotic Translation Initiation Factor 2B (eIF2B).^11^ Disruption of eIF2B activity results in chronic activation of the Integrated Stress Response (ISR).^12,13^ Evidence in both patients and mouse models indicates that astrocyte dysfunction is an important factor in the neurological decline in VWM^14–17^, suggesting that this cell type is a candidate target in the development of a gene therapy for VWM.

In order to investigate the potential of gene addition as a therapy for VWM, we used a lentiviral approach to supplement a defective *Eif2b5^R191H^* variant with the wild-type *Eif2b5* gene (wt*Eif2b5*). A single dose of the therapy was administered in neonates of a VWM mouse model. We tested wt*Eif2b5* overexpression driven by two different promoters: a GFAP promoter to target astrocytes and a CHOP promoter to target cells with increased ISR activation. We studied the effects of wt*Eif2b5* overexpression in adulthood by testing various aspects of the VWM mouse phenotype and provide evidence that by supplementing wt*Eif2b5* in astrocytes, VWM mice show significant improvements in brain pathology and motor skills.

## Results

### Overexpression of wtEif2b5 in astrocytes recovers the oligodendrocyte progenitor cell maturation defect in co-cultures

To test whether overexpression of the wt*Eif2b5* gene in astrocytes can rescue the VWM phenotype, its effects on VWM oligodendrocyte progenitor cell (OPC) maturation were studied. Earlier studies showed that an OPC maturation defect in VWM can be modeled *in vitro* by co-culturing VWM astrocytes with wt OPCs.^16^ Prior to co-culturing, wt (**Fig. 1A**) or VWM astrocytes (**Fig. 1B**) were transduced with lentiviral particles encoding eGFP (‘CMV-eGFP’) and VWM astrocytes were transduced with eGFP-tagged wt*Eif2b5* (‘CMV-wt*Eif2b5*’, **Fig. 1C,D**), driven by the ubiquitous CMV promoter. OPCs co-cultured with CMV-wt*Eif2b5* VWM astrocytes showed stimulated maturation compared to VWM astrocytes transduced with CMV-eGFP (*t*(4)=2.92, *p*=.043, **Fig. 1E**). Indeed, overexpression resulted in levels of myelin basic protein (MBP) expressing cells that were restored to those of OPCs co-cultured with control (*Eif2b5^R191H^* heterozygous) astrocytes (normalized to 1, **Fig. 1A-E**). The eGFP signal in cells treated with CMV-eGFP construct was roughly three times higher than that in cells treated with CMV-eGFP-wt*Eif2b5*, although the intensity of eGFP signal per transduced astrocyte varied widely within both constructs and did not differ statistically from one another(**Fig. 1A-D,F**). These data show that supplementation of wt*Eif2b5* in the VWM genome can alleviate the *in vitro* VWM phenotype of perturbed OPC maturation.

**Figure 1.**
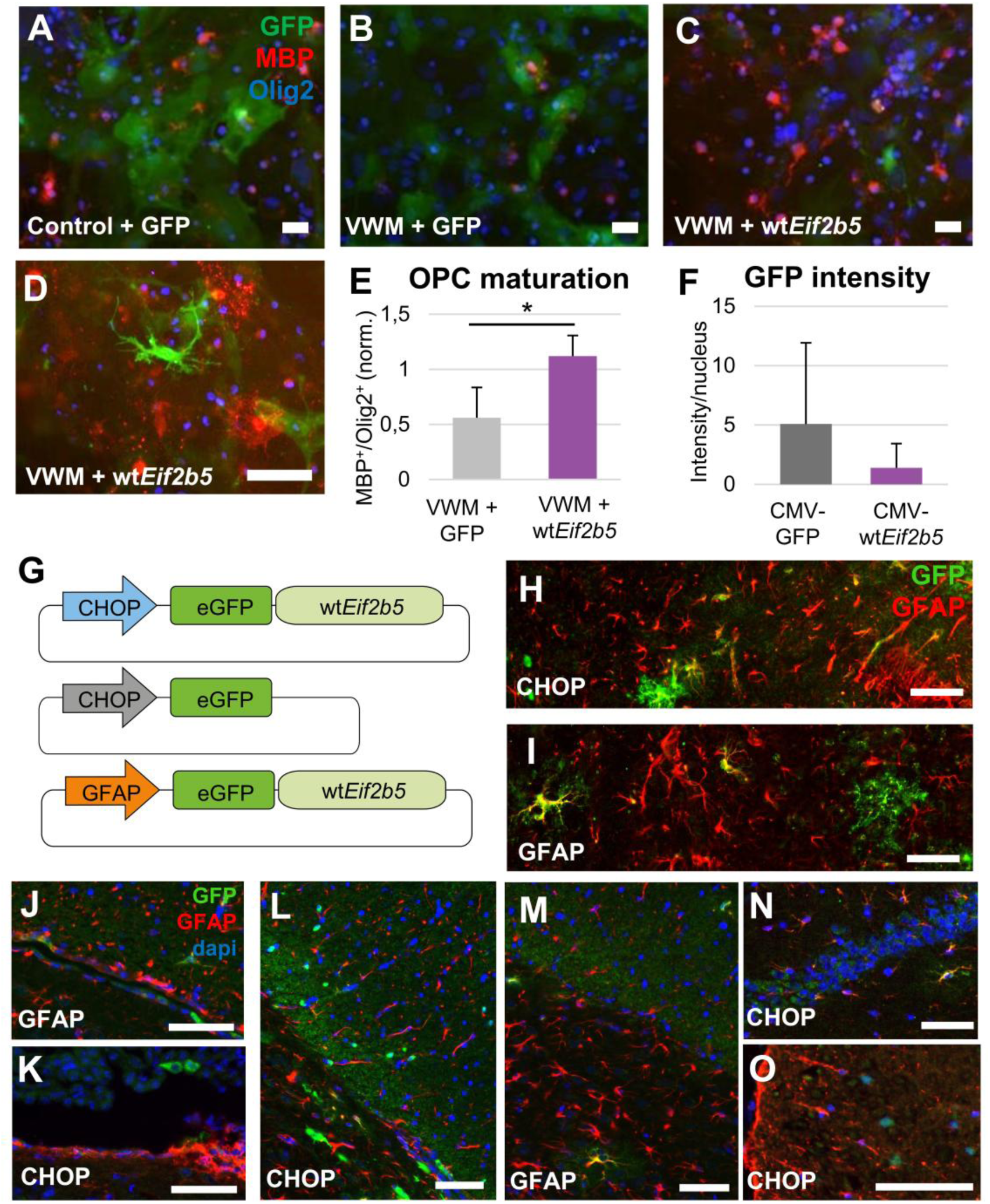
Lentiviral overexpression of wt*Eif2b5* in astrocytes rescues a VWM-induced OPC maturation defect *in vitro* and is expressed in the CNS upon intracerebroventricular administration *in vivo*. Primary co-cultures of wild-type OPCs together with wild-type (control; **A**) or VWM (**B**) astrocytes transduced with CMV-eGFP, or together with VWM astrocytes transduced with CMV-eGFP-wt*Eif2b5* (**C,D**). Quantification of the number of MBP^+^Olig2^+^ oligodendrocytes in the co-culture with wt*Eif2b5*-transduced VWM astrocytes shows levels close to those co-cultured with wild-type astrocytes (normalized to 1; **E**). eGFP expression levels in primary astrocytes after CMV-eGFP and CMV-eGFP-wt*Eif2b5* transduction, as measured by pixel intensity divided by the number of nuclei (**F**). Graphs **E** and **F** represent *n*=4 biological replicates. Schematic overview of the constructs used for *in vivo* lentivirus-mediated overexpression in mouse neonates (**G**). Co-expression of GFAP (red) and CHOP-(eGFP-)wt*Eif2b5* (green; **H**) and of GFAP (red) and GFAP-(eGFP-)wt*Eif2b5* (green; **I**). GFP^+^ cells were observed in the lateral ventricle (**J**), fourth ventricle (**K**), corpus callosum (**L,M**), hippocampus (**N**), and the white matter of the spinal cord (**O**) at 9 months of age following treatment at birth. Independent T-tests were used in **E** and **F**. A one-way ANOVA was used in **G**. *: p-value <.05. All scale bars represent 100µm.

### Supplementation of wtEif2b5 in Eif2b5-mutant VWM mice

To determine whether the beneficial effect of overexpressing wt*Eif2b5* using lentiviral vectors was also observed *in vivo* in *Eif2b5*-mutant VWM mice^16^, we tested the widely used promoters CMV, PGK, and Ubi-C to drive eGFP-tagged wt*Eif2b5* expression (**Fig. S1A**). VWM mice received intracerebroventricular injections of the lentivirus at the day of birth leading to GFP^+^ cells in the brain (**Fig. S1B,E-G**) and the spinal cord (**Fig**. **S1D**). Others showed that these promoters can be active in a variety of cell types (e.g. ^32,33^). The promotors resulted in mostly neuronal tropism in our animals as indicated by co-localization of eGFP with Neuronal Nuclei (NeuN)-labeled cells (**Fig. S1B,C**) and morphology (**Fig. S1E-G)**, and in extremely limited co-labeling of eGFP with glial-associated markers GFAP or Olig2 (data not shown). This pattern was preserved in animals of 5 weeks old, 2 months old, and 9 months old (data not shown). Thus, injection of VSV-G lentiviral particles using the ubiquitous promoters CMV, PGK, and Ubi-C presented a highly neuronal expression of eGFP-tagged wt*Eif2b5 in vivo* when injected at P0.

To test whether specific targeting of cell types affected by VWM would improve the VWM phenotype, the wt*Eif2b5* coding sequence was cloned into constructs including a rodent CHOP or GFAP promoter (**Fig. 1G**). As VWM astrocytes and VWM oligodendrocytes show an increased ISR activation leading to enhanced CHOP expression^12,13^, we expected to specifically target VWM-affected cells by using the CHOP promoter. As astrocytes are considered to be drivers of VWM pathology^16^, we also tested the use of a GFAP promoter to achieve a therapeutic effect. The CNS of animals treated with CHOP-wt*Eif2b5* and GFAP-wt*Eif2b5* showed GFP^+^ cells in the lateral ventricle (**Fig. 1J**), the fourth ventricle (**Fig. 1K**), the corpus callosum (**Fig. 1L,M**), the hippocampus (**Fig. 1N**), and the spinal cord (**Fig. 1O**), amongst other regions.

### Phenotypic improvements in GFAP-wtEif2b5-treated animals

To monitor potential developmental deviations in VWM mice treated with a therapeutic lentivirus vector, animals were weighed weekly. As reported previously, VWM mice have lower body weight than heterozygous controls.^16^ Weight development of treated animals was therefore monitored at 4 months of age onwards, which is when untreated VWM animals start to show a decreased weight.^16^ VWM animals indeed showed lower body weight at 7 months of age, irrespective of treatment (females (*H*(5)=22.50, *p*<.0005, **Fig. 2A**; males (*F*(5,21.4)=24.00, *p*<.0005; **Fig. 2B**). Interestingly, in the group of male VWM animals, the GFAP-wt*Eif2b5* mice had a significantly lower weight than both the VWM CHOP-wt*Eif2b5* (Games-Howell *p*=.032) and GFP controls (Games-Howell *p*=.037; **Fig. 2B**). No such effect was observed in females (**Fig. 2A**). However, no detrimental effects of wt*Eif2b5* overexpression on appearance, motility, feeding, or grooming were observed in any of the groups. Thus, using the GFAP or CHOP promoter, cells implicated in VWM pathophysiology could be targeted for wt*Eif2b5* overexpression without negative effects on weight development or behavior.

**Figure 2.**
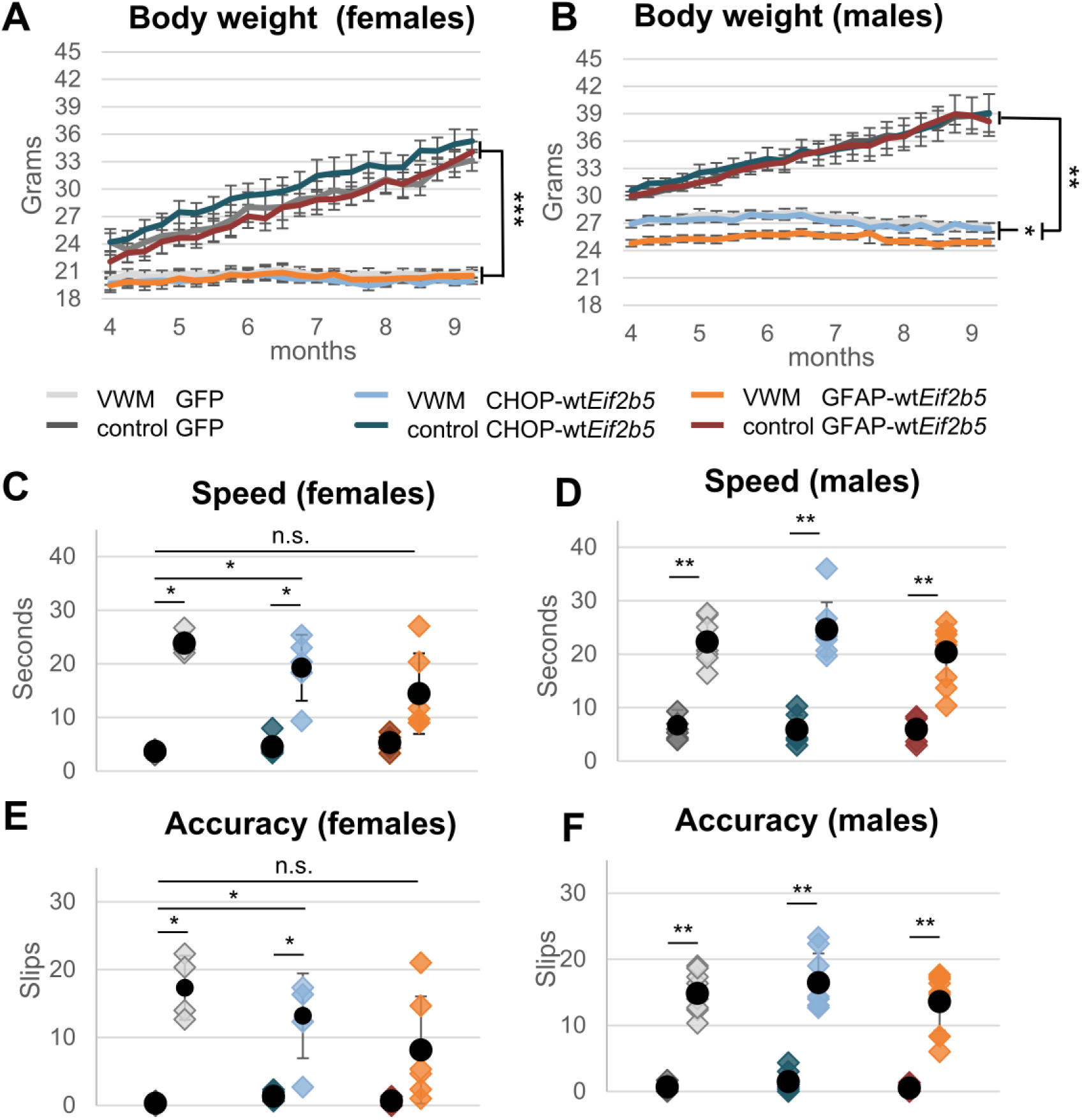
Assessment of effects of GFAP– or CHOP-driven wtEif2b5 supplementation in vivo on the phenotype of Eif2b5-mutant VWM mice. Treatment with CHOP-eGFP (VWM *n*=5, control *n*=3), CHOP-wt*Eif2b5* (VWM *n*=6, control *n*=7), or GFAP-wt*Eif2b5* (VWM *n*=4, control *n*=6) in female mice did not affect the decrease in body weight observed in VWM between 4 to 9 months post-treatment (**A**). A similar observation was made for male animals, although GFAP-wt*Eif2b5* VWM mice (VWM *n*=11, control *n*=8) additionally weighed significantly less than CHOP-eGFP (VWM *n*=9, control *n*=11) and CHOP-wt*Eif2b5* VWM mice (VWM *n*=7, control *n*=8; **B**). At 7 months following treatment at birth, speed (**C, D**) and accuracy (**E, F**) of traversing a narrow beam showed a subset of improved females (**C, E**) and males (**D, F**) in the group that received GFAP-wtEif2b5 treatment. The number of animals per group are listed in Table 1. Kruskal-Wallis tests were used for **A**, **C**, **E** and **F**. A one-way ANOVA with Tukey post-hoc was used in **B** and **D**. *: p-value <.05; **: p-value <.01; ***: p-value <.001; n.s.: non-significant.

**Table 1.** Overview of the number of animals used per treatment, genotype, and sex.

| Treatment | *Eif2b5*<sup>R191H</sup> genotype | Sex |
| --- | --- | --- |
| Saline | Heterozygous | males n=11<br>females n=11 |
|  | Homozygous | males n=6<br>females n=12 |
| CMV-wtEif2b5<br>Titer: $1.46 \times 10^8$ vg/mL | Homozygous | males n=1<br>females n=2 |
| PGK-wtEif2b5<br>Titer: $1.51 \times 10^8$ vg/mL | Homozygous | males n=2<br>females n=3 |
| UBI-wtEif2b5<br>Titer: $1.36 \times 10^8$ vg/mL | Homozygous | males n=1<br>females n=1 |
| Ubiquitous promoters<br>pooled (CMV, PGK, UBI) | Heterozygous | males n=1<br>females n=3 |
|  | Homozygous | males n=4<br>females n=6 |
| CHOP-wtEif2b5<br>Titer: $1.58 \times 10^8$ vg/mL | Heterozygous | males n=8<br>females n=7 |
|  | Homozygous | males n=7<br>females n=6 |
| GFAP-wtEif2b5<br>Titer: $1.47 \times 10^8$ vg/mL | Heterozygous | males n=8<br>females n=6 |
|  | Homozygous | males n=11<br>females n=4 |
| CHOP-eGFP<br>Titer: $1.65 \times 10^8$ vg/mL | Heterozygous | males n=11<br>females n=3 |
|  | Homozygous | males n=9<br>females n=5 |
| CMV-eGFP titer: $1.51 \times 10^8$ vg/mL. | | |

To assess effects of wt*Eif2b5* overexpression on the VWM phenotype, the motor skills of mice that had received wt*Eif2b5* overexpression were tested at 7 months of age. These assays included the traversing of a narrow beam, gripping of a grid, and gait assessment, as described previously.^16^ The CHOP– and GFAP-wt*Eif2b5* overexpression groups showed no clear improvement in the VWM phenotype in terms of stride length (females: *F*(5,7.86)=13.33, *p*=.001, **Fig. S2A**; males: *F*(5,47)=20.27, *p*<.0005; **Fig. S2B**) and step width (females: Kruskal-Wallis test non-significant, **Fig. S2C**; males: *F*(5,47)=20.4.78, *p*=.001, **Fig. S2D**) of the gait assessment, nor in grip strength (females: *F*(5,25)=5.53, *p*=.001, **Fig. S2E**; males: Welch F test non-significant; **Fig. S2F**).

However, half of the VWM animals treated with GFAP-wt*Eif2b5* (n=7) performed better on traversing the balance beam. Particularly in the female GFAP-wt*Eif2b5* group, VWM animals no longer performed worse than healthy animals overexpressing either GFP or GFAP-wt*Eif2b5* in both speed (*H*(5)=21.82, *p*=.001; **Fig. 2C**) and accuracy (*H*(5)=21.13, *p*=.001; **Fig. 2E**). Despite the improved performance of three animals, the male GFAP-wt*Eif2b5* group was still outperformed by healthy controls on the balance beam in terms of speed (*F*(5,43)=45.71, *p*<.0005; **Fig. 2D**) and accuracy (*H*(5)=39.80, *p*<.0005; **Fig. 2F**). One additional GFAP-wt*Eif2b5* male had improved only on step width. For females treated with GFAP-wt*Eif2b5*, one out of the 5 animals that had improved on both balance beam parameters did not show improvement on the step width assay.

VWM animals treated with CHOP-wt*Eif2b5* overall did not show improved motor skills compared to healthy controls, irrespective of sex. However, three animals of the CHOP-wt*Eif2b5* VWM group had improved, with *n*=2 females scoring better on balance beam latency and speed, and *n*=1 male having improved on step width. On balance beam performance, one of the improved females behaved like a positive outlier, performing similar to GFP-overexpressing healthy controls (**Fig. 2C,E**). Importantly, GFAP-wt*Eif2b5* overexpression improved the motor behavior of VWM animals towards control levels.

### Assessment of brain and spinal cord pathology

To further assess the beneficial effects of wt*Eif2b5* overexpression on well-characterized hallmarks of VWM pathology, we analyzed the mice that had improved on the motor tests (from here on referred to as ‘GFAP improved’ or ‘CHOP improved’). CNS tissue of these animals was tested on several VWM disease hallmarks: translocation of Bergmann glia (BG) in the cerebellum^16,34^; Nestin^+^ astrocytes in the corpus callosum in the brain white matter^16,31^; and increased density of unidentified cells in the spinal cord white matter that express, amongst others, Olig2 and Sox9.^26^

Improved animals treated with CHOP-wt*Eif2b5* showed BG localization (*F*(2,13)=9.254, *p*=.003, **Fig. S3A**) and Nestin^+^ levels in the corpus callosum (H(2)=10.637, *p*=.005; **Fig. S3B**) that were comparable to control animals. In the spinal cord white matter, a previously reported^26^ unidentified (dapi^+^) cell population was confirmed in our cohort of VWM animals (*F*(3,12)=11.882, *p*<.001) irrespective of treatment or improvement on motor tests. The higher number of Sox9^+^ cells located within this population in VWM animals was also unchanged by treatment or motor scores (*F*(3,12)=7.3691, *p*<.0005), whereas the increased number of Olig2^+^ cells was decreased only in CHOP-wt*Eif2b5*-treated animals that also showed improvement on motor assays (*F*(3,12)=9.6326, *p*=.002).

To study whether GFAP-wt*Eif2b5* overexpression recovered BG translocation in the cerebellum^16,31^, GFAP improved animals (*n*=7) were compared to GFP-expressing animals. The GFAP improved animals showed significantly reduced BG translocation (**Fig. 3A-C**) and did not statistically differ from GFP-expressing healthy controls (*F*(2,17)=7.914, *p*=.004; **Fig. 3G**). The number of Nestin^+^ astrocytes in the corpus callosum^16^ had also decreased compared to GFP-overexpressing VWM animals (*F*(2,6.74)=75.240, *p*<.0005; **Fig. 3D-F,H**). These data indicate that GFAP-wt*Eif2b5* animals with increased motor skills had also improved on VWM pathology in the brain.

**Figure 3.**
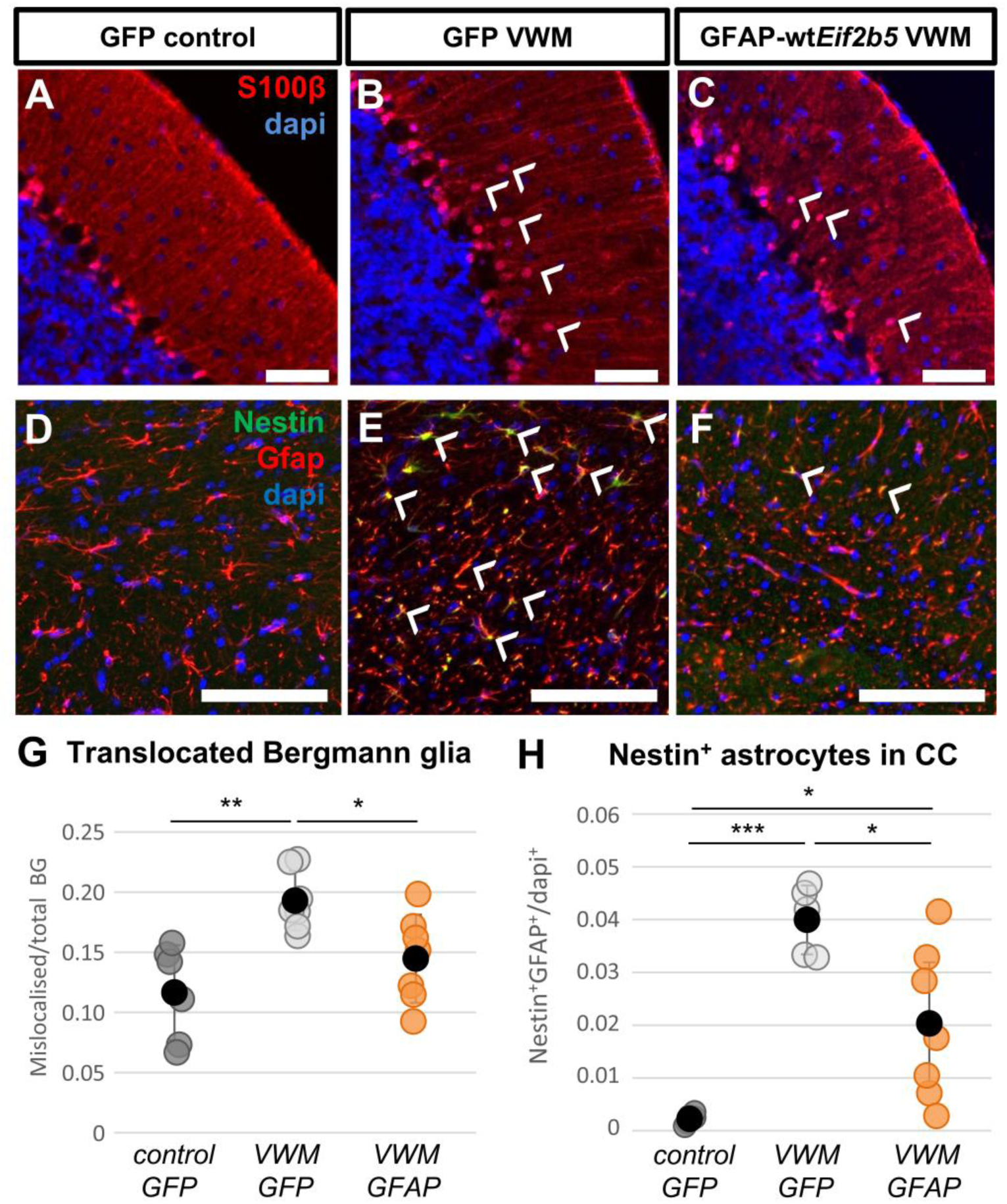
Hallmarks of VWM pathology in the brain were improved after GFAP-wt*Eif2b5* treatment. Animals treated with GFAP-wt*Eif2b5* that had improved on motor tests (*n*=7) showed normalized levels of Bergmann glia translocation as compared to eGFP-expressing control (*n*=6) or VWM animals (*n*=7; arrows, **A-C,G**) at 9 months post-treatment. The number of Nestin^+^ astrocytes in the corpus callosum was also significantly decreased in GFAP-wt*Eif2b5* improved animals (*n*=7) as compared to VWM mice treated with eGFP-only lentivirus (*n*=5; controls *n*=5; **D-F,H**). A one-way ANOVA with Tukey post-hoc was used in **G**. A Welch test with Games-Howell post-hoc was used in **H**. *: p-value <.05; **: p-value <.01; ***: p-value <.001. Scale bars represent 100µm.

In the spinal cord white matter, the dense population of cells in VWM animals (*F*(2,12)=26.658, *p*<.0005) was unaffected by the treatment in the GFAP improved group (**Fig. 4A-C,J**). In contrast, the density of Sox9^+^ cells within the GFAP improved group had decreased to that of healthy GFP controls (*F*(2,7.57=19.804, *p*=.001; **Fig. 4D-F,K**). The density of Olig2^+^ cells had also improved (*F*(2,12)=10.881, *p*=.002; **Fig. 4G-I,L**). Thus, although the overall population density was unaffected, expression of glial lineage markers had normalized to varying extent. These findings indicate that GFAP-wt*Eif2b5* overexpression exerts beneficial effects both on motor skills and on VWM pathology in the spinal cord white matter.

**Figure 4.**
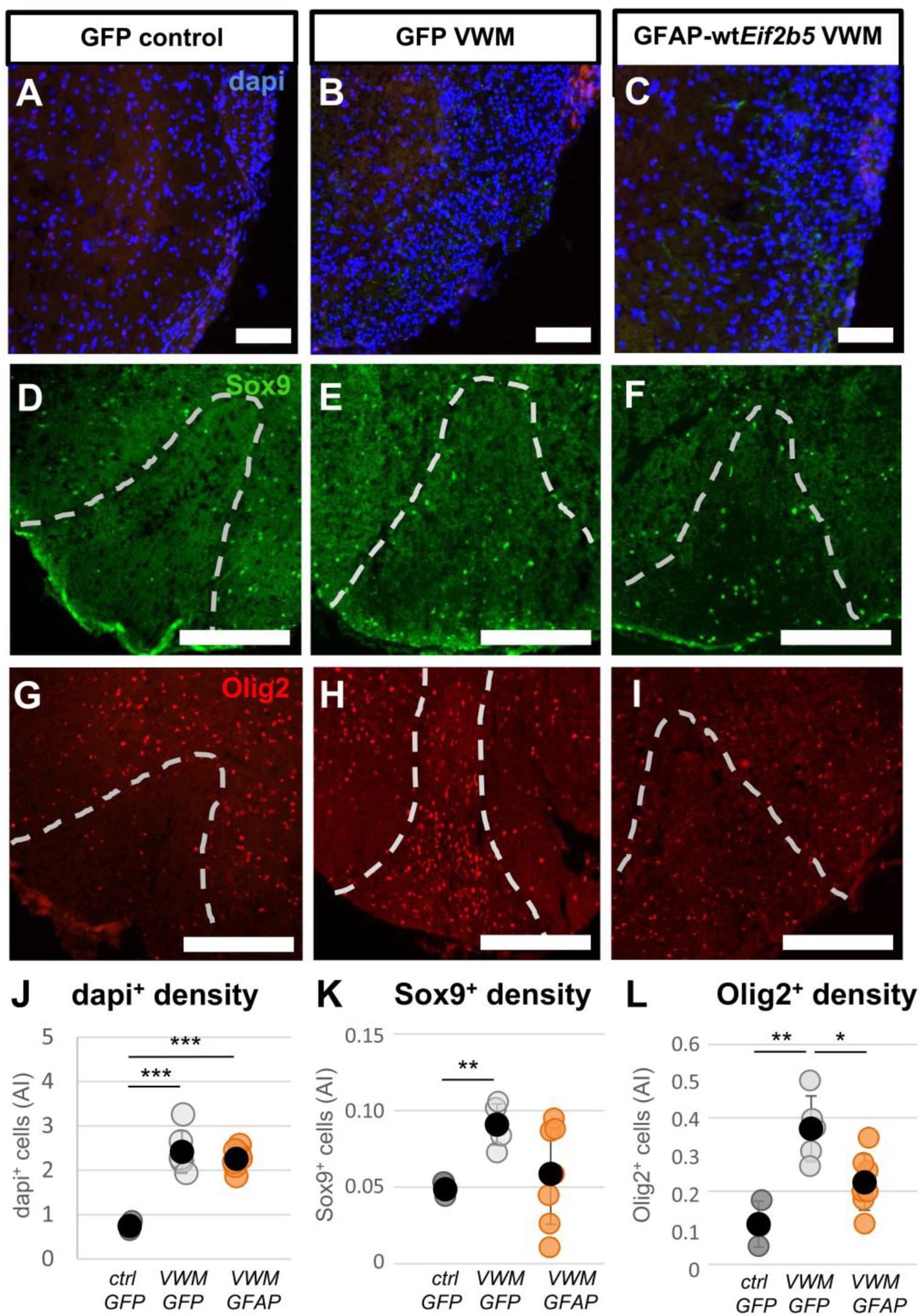
GFAP-wtEif2b5 treatment improved pathological expression of Sox9 and Olig2 in the spinal cord. Spinal cord pathology in eGFP-expressing control (*n*=3) and VWM mice (*n*=5) was compared to GFAP-wt*Eif2b5* VWM mice that had improved on motor assays (*n*=7). At 9 months post-treatment, overall cell density in the white matter was unaffected by treatment (**A-C,J**) whereas the amount of Sox9^+^ cells of this population (**D-F,K**) and the amount of Olig2^+^ cells within this population (**G-I,L**) were improved in the subset of improved GFAP-wtEif2b5 treated mice when analyzed on an arbitrary index (AI) of (number of cells x 1000)/pixel. Spinal cord white matter is demarcated by dashed white lines. A one-way ANOVA with Tukey post-hoc was used in **J** and **L**. A Welch test with Games-Howell post-hoc was used in **K**. *: p-value <.05; **: p-value <.01; ***: p-value <.001. Scale bars represent 100µm.

### Heightened sensitivity of white matter brain pathology to intracerebroventricular GFAP-wtEif2b5 treatment

To study whether the intracerebroventricular administration route more successfully treated the disease parameters in the brain than in the spinal cord, we compared the effects on VWM hallmarks of disease in the brain to the spinal cord in all animals treated with GFAP-wt*Eif2b5*. Compared to GFP-expressing VWM animals, scores of the overall GFAP-wt*Eif2b5* group did not indicate statistically significant improvements in BG translocation or spinal cord pathology (**Fig. 5A-D,F**). The amount of Nestin^+^ astrocytes in the corpus callosum remained significantly decreased (*T*(18)=3.022, *p*=.007; **Fig. 5E,F**). Thus, compared to the spinal cord, astrocyte-related pathology in the brain appeared most sensitive to the therapeutic effects of intracerebroventricular administration of GFAP-wt*Eif2b5* overexpression.

**Figure 5.**
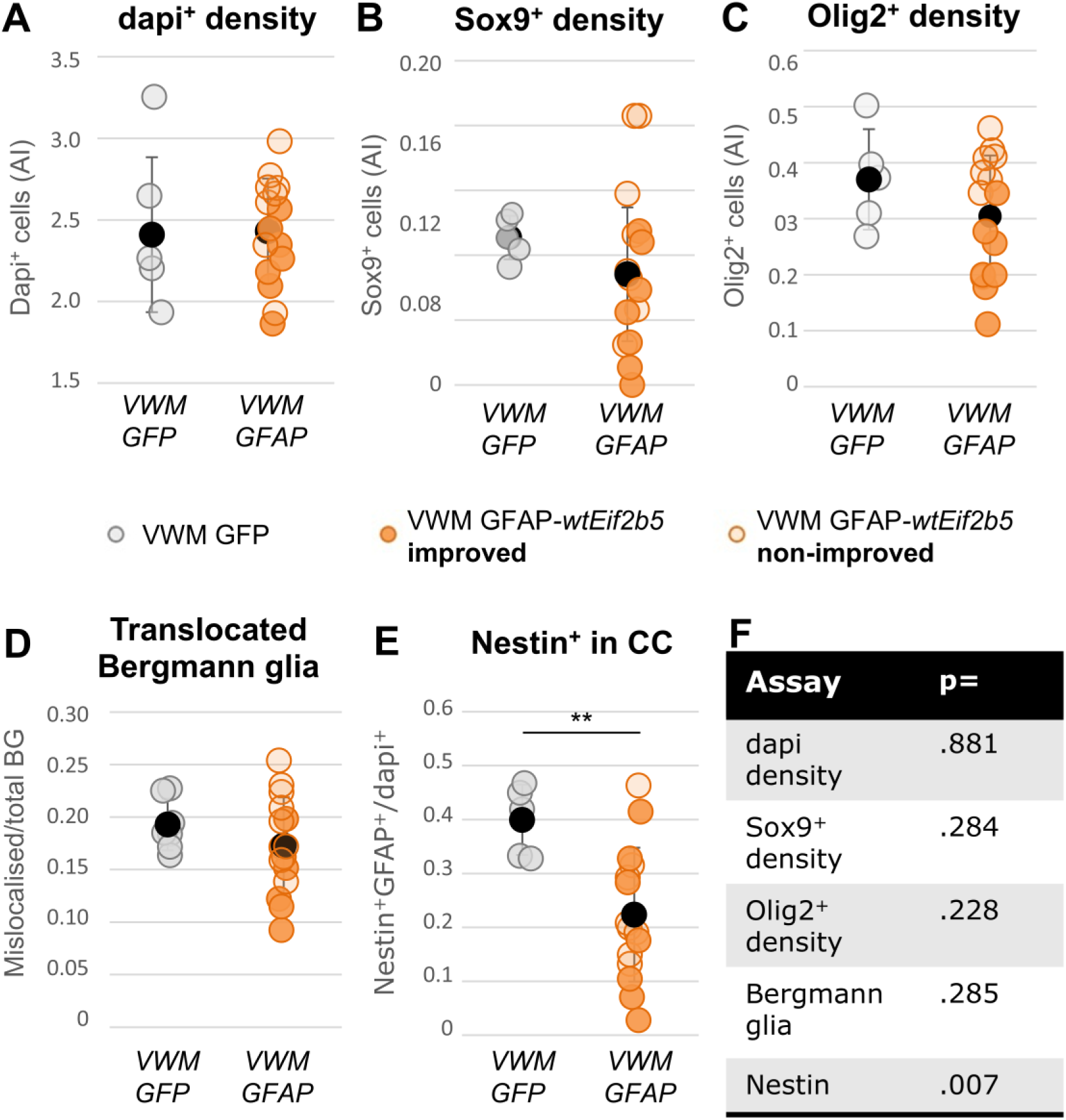
Nestin pathology in brain white matter astrocytes is improved by GFAP-wtEif2b5 treatment. Improved (*n*=7, dark orange) and non-improved (*n*=8, light orange) GFAP-wt*Eif2b5* animals were pooled and compared to CHOP-eGFP VWM animals (*n*=5) on overall cell density (**A**), Sox9^+^ cell density (**B**), Olig2^+^ cell density (**C**) in the spinal cord white matter using an arbitrary index (AI) of (number of cells x 1000)/pixel. Bergmann glia translocation (**D**) and Nestin^+^GFAP^+^ cells (**E**) in the brain of GFAP-wt*Eif2b5* animals (*n*= 15) were compared to VWM animals receiving control treatment (*n*=5) at 9 months post-treatment. The statistical outcome of each comparison is summarized in (**F**). Independent T-tests were used for **A-C** and **E**. A Whitney-Mann U test was performed for **D**. **: p-value <.01.

## Discussion

The current study aimed to develop overexpression-based gene therapy as treatment for the leukodystrophy VWM. Our data shows that the VWM phenotype can be rescued in *Eif2b5^R191H^* mutant mice following intracerebroventricular injection of lentiviral particles encoding the wt *Eif2b5* gene. Motor skills and brain pathology were significantly improved when wt *Eif2b5* overexpression was targeted to astrocytes. Importantly, on case level, half of these treated mice showed improvements in nearly all VWM hallmarks of disease.

As an earlier studied showed that astrocytes are primary affected in VWM^16^ and glial cells display heightened ISR activity^12,13,35^, gene correction in astrocytes was expected to provide strongest therapeutic effects. The precise effects of overexpression remain unexplored on a molecular level, including the localization of idle eIF2Bε subunits and potential clearance mechanisms.

We found that the astrocyte-targeted approach, using GFAP-driven wt*Eif2b5* overexpression, had beneficial effects on the VWM phenotype. Hallmarks of VWM disease, as measured by motor behavior tests and histochemical pathology markers in brain and spinal cord, were improved in half of the treated VWM mice. In line with the astrocytic specificity of GFAP promoter-regulated overexpression, astrocyte-related pathology benefited most from wt*Eif2b5* overexpression. By contrast, only 3 (out of 15) VWM animals treated with CHOP-driven wt*Eif2b5* overexpression showed moderate improvements on motor skills. Pathology in the brain and spinal cord of these animals was partially restored. However, the lack of strong therapeutic effects on motor behavior discouraged further investigation of the CHOP-wt*Eif2b5* vector. In conclusion, wt*Eif2b5* expression regulated by the GFAP promoter reduced pathology severity in the brain and improved motor skills at a systemic level. In-depth understanding of the molecular mechanisms of overexpression in VWM are of interest, considering the dysregulation of protein translation machinery in VWM (e.g. Liu *et al*. ^38^).

The improvements in white matter astrocyte-related pathology are promoted by our use of an astrocyte-specific promoter and the intracerebroventricular route of administration, establishing intracellular wt*Eif2b5* supplementation directly in astrocytes, primarily in the brain. Only this specific targeting of gene supplementation to astrocytes extended therapeutic effects on the molecular pathology to an amelioration of the motor phenotype. However, clinical improvement in motor deficits and spinal cord pathology remained modest whereas pathology in the brain appeared more strongly improved. It remains undetermined whether this is due to the route of administration or because VWM brain pathology is more sensitive to therapeutic intervention. Follow-up studies using intrathecal administration of lentiviral constructs could help determine whether direct targeting of the spinal cord enhances the therapeutic effects on motor deficits.

The therapeutic effects of overexpression using other promoters still need testing. As other studies suggest that oligodendrocytes could contribute to VWM^13^, targeting a common progenitor cell population shared by astrocytes and oligodendrocytes alike^39^ could be of interest. Another option would be to specifically target astrocytes in the white matter, as these have been more strongly implicated in VWM pathology than their grey matter counterparts.^17,40^ Unfortunately, no singular marker exclusive to the white matter astrocyte has been described to our knowledge. The intricate regulation of promoter activity during development, as well as species-dependent differences in cell type specificity of promoters, should not be overlooked either. In specific, our use of the mouse GFAP promoter in neonates ensured a strictly astrocytic expression; by contrast, the human GFAP promoter also targets radial glia^41^ and showed limited expression in the hippocampus and cerebellum^42^ when used in mice. Finally, as diagnosis is often delayed in VWM patients, treatments at later stages are also worthwhile to investigate. In conclusion, investigation of additional promoters and ages of administration could further increase the therapeutic benefits observed by GFAP promoter-regulated wt *Eif2b5* overexpression.

Moreover, a better understanding of the factors driving variability of therapeutic effects across individuals would further contribute to the translational potential of this proof-of-concept VWM gene therapy. Indeed, the varying degree of responsivity of animals to the GFAP-wt*Eif2b5* treatment is of interest for examination. More female than male mice improved after treatment with GFAP-wt*Eif2b5*, indicating that hormonal changes or other sex differences might influence therapeutic effects. Although VWM patients of different sexes show no marked differences in phenotype, ovarian failure is observed in some female VWM patients while no gonadal involvement has been reported in male VWM patients^2,43^. The administration of lentiviral vectors may have also been a source of variation, since stereotaxic coordinates are less reliable when used in neonates.

One limitation of this study is that the original sample collection methodology was not optimal for the assessment of vector copy number per cell. Consequently, our results from this assay are not sufficiently reliable to report. The varying degree of eGFP intensity per cell described in our *in vitro* co-culturing assays suggest that transgene expression *in vivo* is likely also variable. At 9 months of age, the rather limited levels of eGFP detected with immunofluorescence indicate that either initial transduction efficiency was low, yet still sufficient to induce phenotypical improvements, or that expression has decreased over time through silencing of the promoters. Alternatively, clearance of surplus eIF2ε may render eGFP levels low, although the underlying mechanisms remain to be investigated. In the case of the CHOP promoter, the increased baseline ISR activity might have been insufficient to result in notable eGFP expression.

Because integration of HIV-1 is semi-random with a preference for transcriptionally active genomic sites^44,45^, the integration of lentiviral vectors is a concern for mutagenesis. However, the LVs developed for the present study are self-inactivating (SIN) vectors that were designed to limit the risk of insertional activation of nearby genes^46,47^. Lentiviral vector integration is therefore not expected to have led to sufficient insertional mutagenesis to comprise a confounding factor in this study. No phenotypic or histological deviations were observed in the animals to prompt further analysis. The potential therapeutic effects of integrating SIN lentiviral vectors were considered to outweigh a mutagenic potential in this study, considering decreased therapeutic efficacy over time has been observed in an AAV-based gene supplementation study targeting glial cells.^48^

In conclusion, the present study shows that astrocytic overexpression of wt *Eif2b5* in *Eif2b5*-mutant mice improved the VWM phenotype, providing a proof-of-concept that gene therapy can be used as curative treatment option for VWM. The findings in this paper mimic those of successful gene therapies for other leukodystrophies.^8,49–51^ Importantly, no negative effects caused by overexpression were observed in body weight and physical appearance. Potential side effects in the domains of cognition and quality of life, which are difficult to assess in mice, remain to be carefully studied. The gene supplementation approach used in this paper could be a stepping stone towards bringing a curative treatment for VWM to the clinic.

## Materials and methods

### Study design

We aimed to establish whether gene supplementation with *Eif2b5* by lentiviral transduction could improve the phenotype of mouse model for the leukodystrophy VWM. As the primary target for gene supplementation were glial cells that continue to proliferate in healthy and diseased conditions, we opted for an integrative LV vector approach to achieve the maximum number of overexpressing cells over time. The use of episomal vectors such as AAVs would result in a decreased vector copy number per cell with each cell cycle.

An in-house *Eif2b5^R191H^Rag2^null^* VWM mouse model was used. The generation and phenotype of this mouse has been extensively characterized in Dooves *et al*. ^16^ and the model been used in various studies.^12,13,26–28^ Heterozygous littermates (*Eif2b5*^het^) were used as healthy controls, whereas homozygous animals (*Eif2b5*^hom^) develop a VWM phenotype.^16^ We used *Rag2^null^ Eif2b5^R191H^* VWM mice in a C57Bl/6J background to better compare our data to that of previously tested therapies for VWM.^19,20^ VWM mice with or without *Rag2^null^* background^18^ do not perform significantly different when tested on VWM phenotypic assays (*Eif2b5^R191H^Rag2^null^* vs. *Eif2b5^R191H^Rag2^wildtype^* mice; data not shown). To expand the colony with homozygous mutant animals, preferably males or otherwise young females with a *Eif2b5^hom^* genotype were mated with *Eif2b5^het^* animals. Previously characterized phenotypic hallmarks of VWM disease were inspected to study the effects of gene supplementation treatment. We hypothesized that supplementation of *Eif2b5* in astrocytes improves VWM brain pathology (Bergmann glia translocation and immature Nestin-expressing astrocytes in the corpus callosum), and (to lesser extent) spinal cord pathology (Olig2– and Sox9-expressing cells) and motor deficits (balance beam performance, grip strength, gait). Minimal to no effect on body weight was expected.

Each animal was considered one experimental unit. The viral vector treatments were randomized between litters; treatment was the same within a litter for practical reasons (i.e. does not require tracking/marking of newborn pups) and to allow comparison of VWM animals and healthy littermates. Litters were randomized for motor assay testing and researchers were blinded to the litter treatment during assessment. Analysis of the brain and spinal cord pathology of wt*Eif2b5*-treated animals was always combined with the analysis of samples of eGFP-expressing VWM animals and healthy controls to allow for blind assessment.

Effect sizes of phenotypic effects in *Eif2b5^R191H^* mice^16^ were used to perform a power analysis. A β of .80 requires between 4 and 12 animals per genotype per treatment group, i.e. for body weight (required *n*=6), grip strength (required *n*=12), balance beam latency (required *n*=4), and balance beam accuracy (required *n*=9). To account for potential loss of pups (of natural causes, e.g. by poor maternal care), we injected *n*=30 animals per group, resulting in roughly *n*=15 animals per genotype. This group size may also compensate for potentially smaller effects of local, single-dose gene supplementation as compared to the phenotype induced in the *Eif2b5^R191H^* mouse model. The criteria for the humane end point for experimental animals were defined prior to onset of the study and involve a) a weight loss of >15% within a week, b) a decrease of >20% of the highest measured body weight of the animal, or c) severe neurological signs such as loss of muscle tone or paralysis. Upon reaching the humane end point, animals would be euthanized by the same method as described below (using transcardial perfusion under anesthesia by tribromoethanol). At 7 months of age, one animal sustained an infection after fighting with littermates and was euthanized upon poor recovery. No other animals required euthanization. No animals were excluded from analysis, except in the case an animal paused for >10 seconds while crossing the balance beam. We and others have observed extended pausing of animals while traversing the beam^13,21–23^, primarily in control animals that do not present motor deficits. This was the case for *n*=11 healthy control animals (*n*=6 saline controls, *n*=2 eGFP controls, *n*=1 GFAP-wt*Eif2b5*, *n*=1 CHOP-wt*Eif2b5*) and *n*=1 VWM mouse (eGFP-treated). No outliers have been removed from the data. For *in vitro* experiments, *n*=4 biological replicates were analyzed. For *in vivo* experiments, a minimum of *n*=3 animals were analyzed.

### Vector design

A plasmid with a CMV promoter was purchased from Genecopoeia (EX-Mm14891-Lv189) that encoded wt mouse *eIF2bε* with an eGFP tag fused to its N-terminal. To create an ENTRY clone, the *Eif2b5* gene encoded by EX-Mm14891-Lv189 was sub-cloned into pEGFP-ENTRrz1 using KpNI and NotI to maintain the N-terminus fusion of eGFP to *Eif2b5*. This ENTRY clone, pEGFP-*Eif2b5*-ENTRrz1, was used in a Gateway reaction with pDESTFUW and pDEST-PGK(pr)LentiFGA2.0 in order to control expression of eGFP-*Eif2b5* by either a ubiquitin or PGK promoter, respectively. The PGK promoter is derived by PCR from the FRT-PGK-gb2-neo-FRT-loxP plasmid. The Ubi promoter was derived from FUW (Addgene clone #14882). The promoter sequences from CHOP and GFAP were obtained from CLX-CHOP-dGFP (Addgene clone #71299) and GFAP-CRE (Addgene clone #24704) respectively, using PCR technique. The promoters were then sub-cloned in pEGFP-Eif2b5-ENTRrz1 using restriction enzyme AgeI and SLICE cloning technique. *Eif2b5* was removed from pCHOP(pr)EGFP-*Eif2b5*-ENTRrz1 using the restriction enzymes HindIII and NotI to create control construct pCHOP(pr)EGFP-ENTRrz1. The ENTRY vectors were used together with pDESTlentiFGA2.0 in a LR Gateway cloning reaction to create the constructs pCHOP(pr)EGFP-lentiFGA2.0, pCHOP(pr)EGFP-*Eif2b5*-lentiFGA2.0 and pGFAP(pr)EGFP-*Eif2b5*-lentiFGA2.0.

### Lentivirus production and titration

VSV-G-pseudotyped lentiviral particles were produced in HEK293T cells (CRL-11268, ATCC^®^) based on Naldini *et al*.^24^ using a second generation system.^25^ Briefly, cells were cultured in HEK medium (DMEM, 10% fetal calf serum, 50U mL^−1^ penicillin and 50mg mL^−1^ streptomycin (pen/strep, 1%), and 100µM non-essential amino acids) and passaged one day before transfection at ∼50% confluence. Polyethylenimine (5µg/µL, 240µg total) was mixed 3:1 with ENV, PACK, and transfer vector DNA (14.25%, 28.5%, and 57.25% of 80µg, respectively) in 5mL PBS. The mixture was incubated at RT for 10’ prior to adding to the culture. After 24 hours, medium was changed to Optimem (Thermo-Fisher) containing 1% pen/strep. The next day, 40 hours post-transfection, conditioned medium was harvested and collected using Amicon spin filters with a 100kDA cut-off (UFC910024, Millipore). The remaining volume was diluted in 1mL PBS and passed through a 0.2µm filter. The filtered solution was stored in aliquots at –80°C until use. Viral particles were not refrozen after thawing.

HEK293T cells were transduced with each vector at a MOI of 5 (based on a control vector with known titer) in Optimem medium supplemented with 7µg/mL polybrene. Genomic DNA was isolated after 48 hours of incubation. Lentiviral titers were measured by qPCR using forward primer 5’-ggacgtccttctgctacgtc-3’ and reverse primer 5’-aagggagatccgactcgtct-3’ to target the WPRE sequence present in each vector. A Quantstudio 5 Real-time PCR system (384 wells; Applied Biosystems) was used with SensiFast Sybr Hi-ROX (Meridian) to amplify and analyze signal. Data was analyzed with Quantstudio 5 software and showed that titers ranged between 1.4×10^8^ – 1.7×10^8^ vg/mL (M = 1.5×10^8^; SD = 0.094×10^8^). Titers are included in **Table 1**.

### Cell culture

Primary mouse astrocytes were isolated from *Eif2b5*^hom^ VWM mice and *Eif2b5*^het^ healthy control littermates as described previously^16^ and grown in astrocyte medium (DMEM/F12 supplemented with 10% FBS, 1% pen/strep, and 1% L-glutamate) in uncoated T75 flasks. Medium was refreshed once to twice a week. Cells were passaged twice to obtain a purer astrocyte population.

### In vitro lentiviral transduction

During the third passage, cell suspensions were inoculated with virus (MOI = 4.8 for CMV-*wt*Eif2b5; MOI = 5.0 for CMV-eGFP) for overnight transfection. The next morning, transduced astrocytes were washed with PBS once and were maintained as described above until confluent. At the next and final passage, transfected astrocytes were additionally spinoculated with lentivirus (MOI = 3.6 for CMV-*wt*Eif2b5; MOI = 3.8 for CMV-eGFP) and polybrene (4µL/mL) before being plated. VWM astrocytes were transfected with either CMV-eGFP-*Eif2b5* or CMV-eGFP virus. Healthy control astrocytes were transfected with CMV-eGFP only.

### OPC-astrocyte co-cultures

One week prior to adding OPCs, transfected astrocytes were plated directly following spinoculation in an 8-well chamber slide (Nunc Lab-tek) at a density of 100.000 cells/well. Cells were washed the day after and full medium was added. P0 wt mouse brains were isolated in HBSS^-^, dissociated with MACS Gentle Dissociator, and incubated at 37°C overnight. The next day, medium of transfected astrocytes was switched to OPC medium.^29^ The dissociated cells were sorted for PDGFαR^+^ OPCs using MACS according to manufacturer’s protocol (CD140a antibody, 130-102-473, Miltenyi). Sorted OPCs were directly plated onto transfected astrocytes at a density of 100.000 cells/well. Co-cultures were kept for one week before being washed with PBS and fixed in 2% paraformaldehyde for 20 minutes. The cultures were washed once more and used for immunocytochemistry.

### Intracerebroventricular (ICV) virus injections in neonatal mice

ICV injections were adapted from Kim *et al*.^30^ with an instrumental set-up described by Dooves *et al*.^31^ Briefly, P0-P1 mice were anesthetized on ice, and were bilaterally injected in the lateral ventricles with a 34G beveled needle (NanoFil). The needle was mounted to a stereotaxic set-up and controlled by an automated pump (World Precision Instruments) to inject 3µL virus into each ventricle (8.1 – 9.9 x 10^5^ viral particles/neonate total) at a rate of 1µL per minute. Coordinates starting from Lambda were AP: +1.5mm, ML: ±0.8mm, DV: –1.7mm to –1.5mm.^30^ The number of animals injected per treatment and the virus titers of respective vectors are listed in **Table 1**, specified per VWM genotype (*Eif2b5^R191H^* heterozygous or homozygous) and sex.

### Motor tests of 7-month-old mice

Animals were tested on a variety of motor skills as previously described by Dooves *et al*.^16^ In short, animals were tested on speed and accuracy of crossing a narrow beam, on grip strength of the front limbs and of all limbs combined, and on gait characteristics such as stride length and step width ratio of the hind and front paws. The grip strength score represents the mean of *n*=5 consecutive trials within one session. The balance beam score is composed of the average of three sessions. Observers were blind to the phenotype and treatment of the mice. Litters were tested on motor assays in a randomized fashion.

### Immunocyto– and immunofluorescence (ICC and IF)

ICC and IHC was performed as described previously.^16,17^ ICC was used to visualize a nuclear OPC marker (Olig2), Myelin Basic Protein (MBP) in matured oligodendrocytes, and the viral eGFP tag in astrocyte-OPC co-cultures. The cultures were washed 3 times 10’ in PBS before adding blocking buffer (PBS supplemented with 5% normal goat serum, 0.3% Triton-X, and 0.1% BSA) for 1hr at room temperature (RT). Cultures were incubated with antibodies against MBP (Covance, 1:2000, SMI-99P), Olig2 (Millipore, 1:1000, AB9610), and GFP (Aves Labs, 1:1000, GFP-1020) overnight at 4°C. Cells were then washed in PBS and incubated with secondary antibodies (Alexa Fluor 594, 647, or 488 respectively, 1:1000, Fisher Scientific) for 1hr at RT. Cultures were washed again before and after counterstaining with DAPI, and were stored at 4°C.

For immunofluorescence, animals were euthanized by intraperitoneal injection with 50mg/kg tribomoethanol. When fully unresponsive, animals were transcardially perfused with 20mL of saline followed by 10mL of 4% PFA. Central nervous system (CNS) tissue was post-fixated in 4% PFA under agitation for 24 hours. Tissue was isolated from the bone structures (skull, spinal column) and incubated under agitation in 30% sucrose in PBS until saturated. Finally, tissue was embedded in Optimal cutting temperature compound (OCT; Sakura Finetec) and frozen in a bath of 2-methylbutane and dry ice. Tissue was stored at –80°C until use. Tissue was cut into 12µm sagittal sections on a cryostat. Brain and spinal cord slices were then used for IF as described previously^16^ to visualize Bergmann glia in the cerebellum using S100β (ProteinTech, 1:1000, 15146-1-AP), Nestin^+^ astrocytes in the corpus callosum (Nestin: BD Bioscience, 1:500, 611658; GFAP: DAKO, 1:1000, Z0334), and cells expressing Sox9 (Cell Signaling, 1:500, 82630) or Olig2 (Millipore, 1:1000, AB9610) in the spinal cord white matter. DAPI was used as a counterstain.

### Microscopy

ICC and IHC was visualized with a Leica DM6000B microscope (Leica Microsystems) using LAS AF software version 2.7.0.9329.

### Cell counts

All counts were performed by a blind observer. Since OPCs tended to cluster at the outer edge of the wells, this entire area of the well was imaged at 200x magnification, as well as 5 images of the center of the well. Olig2– and MBP-positive cells were counted manually using ImageJ software.

Counts of translocated Bergmann glia and Nestin^+^ astrocytes in the corpus callosum were performed as described earlier.^16^ The amount of dapi-positive cells in the white matter of the spinal cord were counted in *n*=3 thoracic sections per animal using the Image-based Tool for Counting Nuclei (ITCN) plug-in for ImageJ with Width set to 12 and Minimum Distance set to 6. Olig2^+^ and Sox9^+^ cells in the white matter of the spinal cord of *n*=3 thoracic sections per animal were assessed as described in Leferink *et al*.^26^, also using the ITCN plug-in.

### Statistical analysis

Improved animals were defined as having a better test score than the best performing eGFP-treated VWM mouse on balance beam speed, latency and/or the ratio of step width of the hind/front paws.

All data is represented as mean ± standard deviation (also see **Supplemental Table 1**). Statistical tests were performed in SPSS version 26.0 (IBM Statistics). All statistical tests were two-sided. A Shapiro-Wilk test was performed to determine normality of the data. When comparing two groups, a Student’s t-test or Whitney-Mann U test was used when appropriate. When comparing more groups, a one-way ANOVA was performed with Tukey post-hoc testing. In case of unequal variances as indicated by a significant Levene’s test, a Welch test with Games-Howell post-hoc testing was performed. If assumptions of a statistical test were not met, non-parametric testing was used. Comparisons of two groups were made using Whitney-Mann U-tests; multiple groups were compared with a Kruskal-Wallis test. The number of animals used for each assay and the associated statistical descriptives are listed in **Supplemental Table 1**.

### Study approval

All experiments were approved by the Animal Welfare Committee of the Vrije University Amsterdam.

### Data availability

All data is available upon reasonable request.

## Acknowledgements

Author contributions are as follows: Conceptualization: A.E.J.H., M.v.d.K, V.M.H.; Data collection: A.E.J.H, R.Z.; Formal Analysis: A.E.J.H; Funding acquisition: M.v.d.K., V.M.H.; Investigation: A.E.J.H.; Methodology: A.E.J.H, R.Z. M.v.d.K., V.M.H.; Resources: M.v.d.K, V.M.H.; Supervision: V.M.H.; Visualization: A.E.J.H.; Writing – original draft: A.E.J.H.; Writing – review & editing: A.E.J.H., M.v.d.K, V.M.H. M.v.d.K. is a member of the European reference network for rare neurological disorders (ERN-RND), project ID 739510. We thank Jeroen Pasterkamp for gifting the PGK promoter sequence, David Baltimore for gifting the Ubi promoter sequence from Addgene clone #14882, Patrick Salmon for the CHOP promoter sequence from Addgene clone #71299, and Michael Sofroniew for the GFAP promoter sequence from Addgene clone #24704.

