## Supplemental material for "Gene supplementation in Vanishing White Matter mice ameliorates the disease"

### Supplementary material

#### Supplemental figures

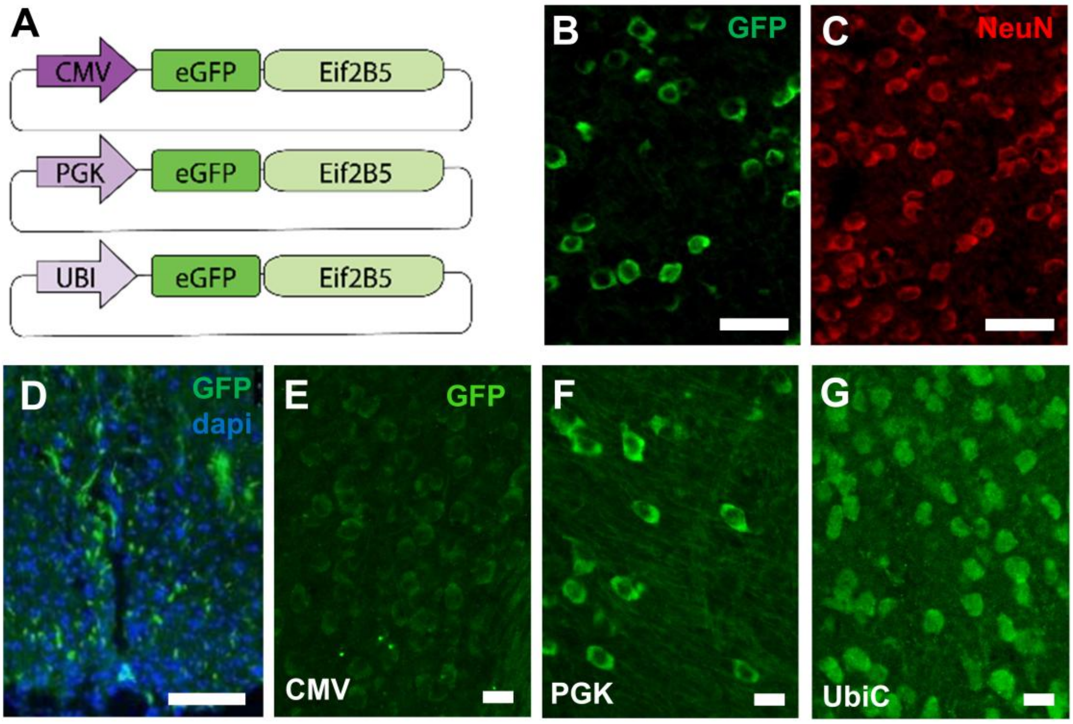


**Supplemental figure 1. Representative imagines of transgene expression in animals injected with eGFP-wt*Eif2b5* under the ubiquitous promoters CMV, PGK, or Ubi-C.** Three vectors with different ubiquitous promoters driving eGFP-wt*Eif2b5* expression in VWM mice were used to generate lentiviral particles (**A**). Following intracerebroventricular injections at P0, eGFP$^+$ cells were found in the cortex (**B,E-G**) and the spinal cord (**D**) at 9 months of age. Most eGFP$^+$ cells showed co-localization with NeuN (**B,C**) and displayed a neuronal morphology (**E-G)**.

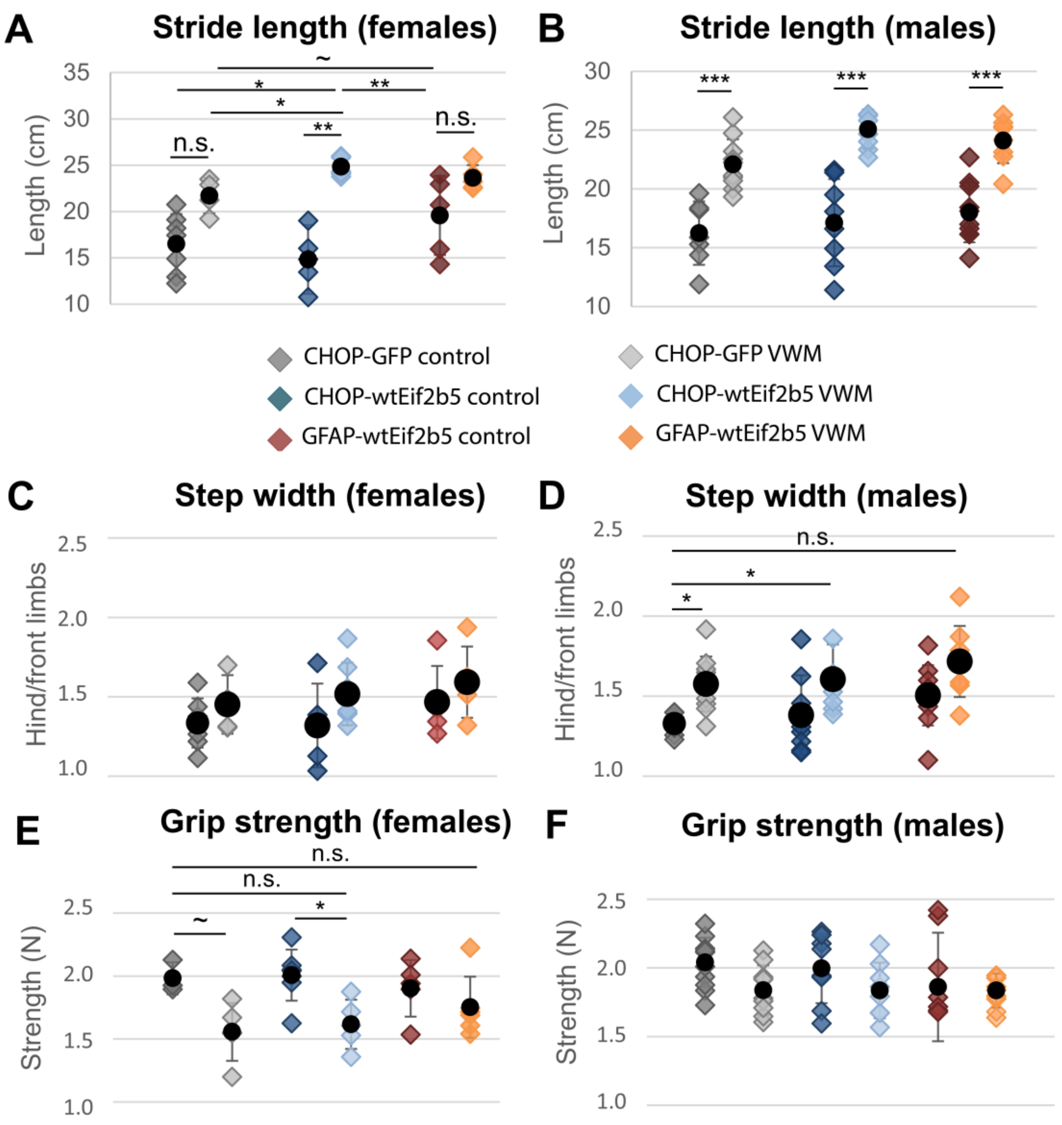


**Supplemental figure 2. Gait assessment and grip strength of animals treated with VWM-specific vectors (CHOP-wt*Eif2b5*, GFAP-wt*Eif2b5*, or CHOP-eGFP).** Stride length (**A**,**B**), ratio of front/hind paw step width (**C**,**D**), and grip strength of front and hind paws (**E**,**F**) are shown for female (**A**,**C**,**E**) and male (**B**,**D**,**F**) mice. The number of mice in each group is indicated in Table 1. A one-way ANOVA with Tukey post-hoc was used for **A**, **D**, and **E**. A Welch test with Games-Howell post-hoc was used for **B** and **E**. A Kruskal-Wallis test was used for **C**. *: $p<.05$; **: $p<.01$; ***: $p<.001$, ~: statistical trend.

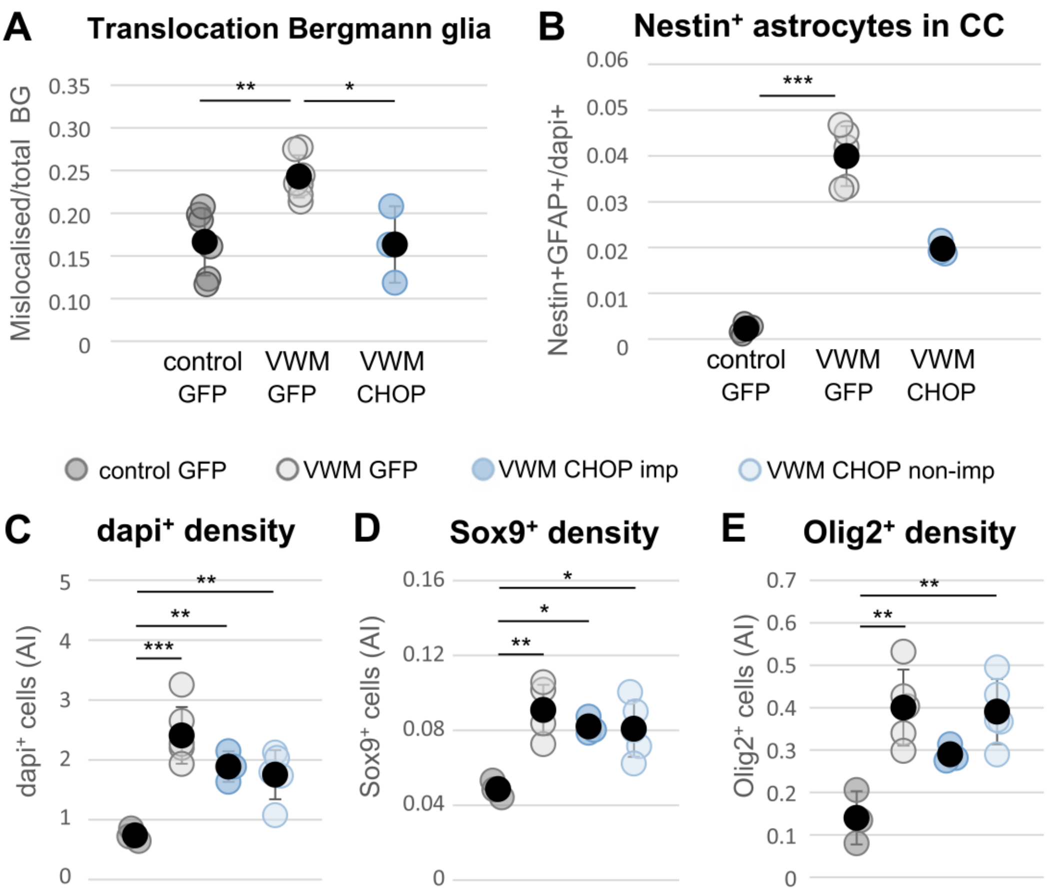


**Supplemental figure 3. Brain and spinal cord pathology following CHOP-wt*Eif2b5* treatment in animals, divided by improvement on motor assays.** Bergmann translocation was normalized when compared to CHOP-eGFP-expressing control (*n*=6) and VWM animals (n=7; **A**). Nestin expression decreased to levels comparable towards that of CHOP-eGFP expressing control animals (control, n=5; VWM, n=5; **B**). Spinal cord white matter pathology after CHOP-wt*Eif2b5* treatment in animals had improved on motor tests ('imp', n=3) or not ('non-imp', n=5) in terms of overall (dapi$^+$) cell density (**C**), Sox9$^+$ cells (**D**), and Olig2$^+$ cells (**E**). A one-way ANOVA with Tukey post-hoc was used for **A, C-E**. A Kruskal-Wallis test was used for **B**. *: *p*<.05; **: *p*<.01; ***: *p*<.001.

**Supplemental Table 1.** Number of animals used and statistical descriptors per assay

| **Experimental group** | **Assay** | ***N*** | **M + SD (or median + range)** | **95% confidence interval** |
|---|---|---|---|---|

| | | | | |
|---|---|---|---|---|
| CHOP-eGFP-wt*Eif2b5* | Body weight at 7 months of age | Females<br>control *n*=7<br>VWM *n*=6<br>Males<br>control *n*= 8<br>VWM *n*= 7 | M=31.16, SD±4.18<br>M=20.97, SD±2.53<br>M=35.38, SD±4.33<br>M=27.27, SD±1.00 | 27.29-35.02<br>18.31-23.62<br>31.76-38.99<br>26.35-28.20 |
| | Balance beam speed | Females<br>control *n*=6<br>VWM *n*=6<br>Males<br>control *n*= 7<br>VWM *n*= 7 | M=4.57, SD±3.05<br>M=22.03, SD±8.78<br>M=5.90, SD±2.65<br>M=23.01, SD±2.31 | 2.73-6.40<br>12.82-31.25<br>3.45-8.35<br>20.88-25.15 |
| | Balance beam accuracy | Females<br>control *n*=7<br>VWM *n*=6<br>Males<br>control *n*= 7<br>VWM *n*= 7 | M=1.19, SD±0.73<br>M=15.15, SD±6.04<br>M=1.47, SD±1.58<br>M=16.09, SD±4.63 | 0.51-1.86<br>7.81-20.49<br>0.01-2.93<br>11.80-20.37 |
| | Grip strength, front+hind | Females<br>control *n*=7<br>VWM *n*=6<br>Males<br>control *n*= 8<br>VWM *n*= 7 | M=2.01, SD±0.21<br>M=1.65, SD±0.19<br>M=1.99, SD±0.25<br>M=1.84, SD±0.22 | 1.82-2.20<br>1.45-1.84<br>1.78-2.21<br>1.64-2.04 |
| | Step width | Females<br>control *n*=7<br>VWM *n*=6<br>Males<br>control *n*= 7<br>VWM *n*= 7 | M=1.52, SD±0.20<br>M=1.29, SD±0.25<br>M=1.61, SD±0.22<br>M=1.42, SD±0.25 | 1.34-1.70<br>1.03-1.55<br>1.41-1.81<br>1.18-1.65 |

| | | | | |
|---|---|---|---|---|
| | Stride length | Females | | |
| | | control *n*=7 | M=24.84, SD±0.99 | 23.92-25.76 |
| | | VWM *n*=6 | M=14.25, SD±3.07 | 11.04-17.47 |
| | | Males | | |
| | | control *n*= 7 | M=24.95, SD±1.52 | 23.54-26.36 |
| | | VWM *n*= 7 | M=17.95, SD±3.12 | 15.06-20.84 |
| | Bergmann glia translocation | VWM imp *n*=3 | M=0.113, SD±0.05 | 0.001-0.225 |
| | $Nestin^{+}GFAP^{+}$ in cc | VWM imp *n*=3 | M=0.0197, SD±0.001 | 0.017-0.023 |
| CHOP-eGFP | Body weight | Females | | |
| | | control *n*= 3 | M=28.73, SD±2.25 | 23.14-34.32 |
| | | VWM *n*= 5 | M=20.32, SD±1.58 | 18.36-22.28 |
| | | Males | | |
| | | control *n*= 11 | M=35.44, SD±5.39 | 31.82-39.06 |
| | | VWM *n*= 9 | M=27.57, SD±1.61 | 26.33-28.80 |
| | Balance beam speed | Females | | |
| | | control *n*=3 | M=3.67, SD±0.58 | 2.23-5.10 |
| | | VWM *n*=4 | M=23.85, SD±2.07 | 20.55-27.15 |
| | | Males | | |
| | | control *n*= 9 | M=6.69, SD±2.91 | 4.45-8.93 |
| | | VWM *n*= 9 | M=22.29, SD±3.78 | 19.38-25.20 |
| | Balance beam accuracy | Females | | |
| | | control *n*= 3 | M=0.33, SD±0.35 | -0.54-1.21 |
| | | VWM *n*= 5 | M=14.52, SD±7.47 | 5.24-23.80 |
| | | Males | | |
| | | control *n*= 11 | M=0.67, SD±0.56 | 0.30-1.05 |
| | | VWM *n*= 9 | M=14.81, SD±3.11 | 12.42-17.20 |
| | Grip strength, front+hind | Females | | |
| | | control *n*= 3 | M=1.98, SD±0.13 | 1.66-2.30 |
| | | VWM *n*= 5 | M=1.56, SD±0.23 | 1.27-1.84 |
| | | Males | | |

| | | | | |
|---|---|---|---|---|
| | | control *n*= 11<br>VWM *n*= 9 | M=2.04, SD±0.18<br>M=1.84, SD±0.18 | 1.92-2.16<br>1.70-1.98 |
| | Step width | Females<br>control *n*= 3<br>VWM *n*= 5<br>Males<br>control *n*= 11<br>VWM *n*= 9 | <br>M=1.44, SD±0.22<br>M=1.37, SD±0.15<br><br>M=1.57, SD±0.16<br>M=1.31, SD±0.10 | <br>0.88-2.00<br>1.19-1.55<br><br>1.46-1.68<br>1.24-1.38 |
| | Stride length | Females<br>control *n*= 3<br>VWM *n*= 5<br>Males<br>control *n*= 11<br>VWM *n*= 9 | <br>M=21.87, SD±5.13<br>M=16.49, SD±3.77<br><br>M=22.00, SD±2.03<br>M=16.30, SD±2.48 | <br>16.24-27.49<br>11.81-21.18<br><br>20.64-23.37<br>14.39-18.20 |
| | Bergmann glia translocation | control *n*= 6<br>VWM *n*= 7 | M=0.12, SD±0.02<br>M=0.19, SD±0.03 | 0.075-0.159<br>0.167-0.218 |
| | Nestin$^+$GFAP$^+$ in cc | control *n*= 5<br>VWM *n*= 5 | M=0.002, SD±0.001<br>M=0.040, SD±0.007 | 0.0006-0.004<br>0.032-0.048 |
| | Sox9$^+$ in s.c. | control *n*= 3<br>VWM *n*= 5 | M=0.048, SD±0.005<br>M=0.912, SD±0.13 | 0.037-0.060<br>0.075-0.108 |
| | Olig2$^+$ in s.c. | control *n*= 3<br>VWM *n*= 5 | M=0.110, SD±0.063<br>M=0.370, SD±0.090 | -0.047-0.267<br>0.259-0.481 |
| | Dapi in s.c. | control *n*= 3<br>VWM *n*= 5 | M=0.743, SD±0.11<br>M=2.460, SD±0.51 | 0.478-1.009<br>1.825-3.095 |
| GFAP-eGFP-wt*Eif2b5* | Body weight | Females<br>control *n*= 6<br>VWM *n*= 4<br>Males<br>control *n*= 8<br>VWM *n*= 11 | <br>M=28.90, SD±3.09<br>M=20.50, SD±1.51<br><br>M=35.55, SD±2.91<br>M=25.15, SD±1.61 | <br>25.66-32.14<br>18.10-20.44<br><br>33.12-37.98<br>24.06-26.23 |

| | | | | |
|---|---|---|---|---|
| | Balance beam speed | Females<br>control *n*= 6<br>VWM *n*= 4<br>Males<br>control *n*= 6<br>VWM *n*= 11 | <br>M=5.08, SD±1.51<br>M=12.68, SD±5.21<br><br>M=5.95, SD±2.19<br>M=20.97, SD±5.35 | <br>3.50-6.67<br>4.38-20.97<br><br>3.65-8.25<br>17.38-24.57 |
| | Balance beam accuracy | Females<br>control *n*= 6<br>VWM *n*= 4<br>Males<br>control *n*= 8<br>VWM *n*= 11 | <br>M=0.93, SD±0.79<br>M=7.25, SD±9.29<br><br>M=0.46, SD±0.53<br>M=13.66, SD±4.12 | <br>0.100-1.767<br>-15.94-34.61<br><br>0.02-0.91<br>10.90-16.43 |
| | Grip strength, front+hind | Females<br>control *n*= 6<br>VWM *n*= 4<br>Males<br>control *n*= 8<br>VWM *n*= 11 | <br>M=1.96, SD±0.24<br>M=1.64, SD±0.08<br><br>M=1.86, SD±0.40<br>M=1.83, SD±0.12 | <br>1.70-2.21<br>1.51-1.76<br><br>1.53-2.19<br>1.75-1.91 |
| | Step width | Females<br>control *n*= 6<br>VWM *n*= 3<br>Males<br>control *n*= 8<br>VWM *n*= 11 | <br>M=1.57, SD±0.21<br>M=1.54, SD±0.27<br><br>M=1.72, SD±0.22<br>M=1.48, SD±0.19 | <br>1.35-1.79<br>0.86-2.22<br><br>1.53-1.90<br>1.35-1.61 |
| | Stride length | Females<br>control *n*= 6<br>VWM *n*= 3<br>Males<br>control *n*= 8<br>VWM *n*= 11 | <br>M=23.73, SD±1.25<br>M=19.86, SD±3.61<br><br>M=24.12, SD±1.90<br>M=17.68, SD±2.66 | <br>22.42-25.04<br>10.89-28.84<br><br>22.53-25.71<br>15.89-19.46 |

| | | | | |
|---|---|---|---|---|
| | Bergmann glia translocation | VWM imp $n$=7<br>VWM (all) $n$=15 | M=0.143, SD±0.04<br>M=0.172, SD±0.46 | 0.108-0.178<br>0.146-0.198 |
| | Nestin$^+$GFAP$^+$ in cc | VWM imp $n$=7<br>VWM (all) $n$=15 | M=0.200, SD±0.14<br>M=0.223, SD±0.12 | 0.007-0.033<br>0.016-0.029 |
| | Sox9$^+$ in s.c. | VWM imp $n$=7<br>VWM (all) $n$=15 | M=0.059, SD±0.03<br>M=0.071, SD±0.04 | 0.028-0.89<br>0.048-0.093 |
| | Olig2$^+$ in s.c. | VWM imp $n$=7<br>VWM (all) $n$=15 | M=0.224, SD±0.08<br>M=0.297, SD±0.12 | 0.153-0.294<br>0.231-0.363 |
| | Dapi in s.c. | VWM imp $n$=7<br>VWM (all) $n$=15 | M=2.25, SD±0.056<br>M=2.43, SD±0.323 | 2.033-2.470<br>2.251-2.610 |
